# Spatial patterns of species richness and nestedness in ant assemblages along an elevational gradient in a Mediterranean mountain range

**DOI:** 10.1101/420059

**Authors:** O. Flores, J. Seoane, V. Hevia, F.M. Azcárate

## Abstract

The study of biodiversity spatial patterns along ecological gradients can serve to elucidate factors shaping biological community structure and predict ecosystem responses to global change. Ant assemblages are particularly interesting as study cases, because ant species play a key role in many ecosystem processes and have frequently been identified as useful bioindicators. Here we analyzed the response of ant species richness and assemblage composition to elevational gradients in Mediterranean grasslands and subsequently tested whether these responses were stable spatially and temporally. We sampled ant assemblages in two years (2014, 2015) in two mountain ranges (Guadarrama, Serrota) in Central Spain, along an elevational gradient ranging from 685 to 2390 m a.s.l.

Jackknife estimates of ant species richness ranged from three to 18.5 species and exhibited a hump-shaped relationship with elevation that peaked at mid range values (1100 - 1400 m). This pattern was transferable temporally and spatially. Elevation was significantly related to ant assemblage composition and facilitated separation of higher elevation assemblages (> 1700 m) from the remaining lower elevation species groups. Ant assemblages were nested; therefore species assemblages with a decreased number of species were a subset of the richer assemblages, although species turnover was more important than pure nestedness in all surveys. The degree of nestedness changed non-linearly as a cubic polynomial with elevation. These assembly patterns were observed over time but not between the two study regions.

We concluded double environmental stressors typical of Mediterranean mountains explained species richness patterns: drought at low elevations and cold temperatures at high elevations likely constrained richness at both extremes of elevational gradients. The fact that species turnover showed a dominant role over pure nestedness suggested current ant assemblages were context-dependent (spatio-temporal factors) and highly vulnerable to global change, which threatens the conservation of present day native ant communities, particularly at high elevations.

## Introduction

Predicting the response of biodiversity to the main drivers of Global Change has become a primary goal of modern ecology [1–3]. Consequently, the analysis of species richness and assembly patterns along latitudinal and altitudinal gradients provide clues to project possible effects of climate change on communities [4, 5]. The Mediterranean mountains are particularly suitable for this purpose, given a characteristic combination of temperature and water availability gradients in space and time, which determines a progressively cooler and wetter environment at higher elevations while valleys suffer more severe summer drought conditions [6, 7]. In addition, the Mediterranean Basin will probable face particularly marked increases in aridity, temperature and frequency of extreme climatic events [8–10]; therefore it is critical to advance our knowledge of how basin communities respond to more severe abiotic factors.

Ants are diverse and ubiquitous, exhibit strong interspecific interactions, and perform many functional processes in ecosystems [11]; therefore ants are considered a viable indicator group of ecosystem change by many community ecologists. Ants are also dominant terrestrial community members [12] and play an important role in plant community dynamics, acting as seed harvesters [13], dispersal agents [14], and influencing soil nutrient status and plant growth [15]. As predators, ants control herbivore abundance, drive relevant top-down effects, and are often used as biological control agents of insect pests and fungal pathogens [16]. Furthermore, ant distribution and abundance patterns provide information to address specific environmental management issues [17, 18]; consequently, ants are particularly useful bioindicators.

Previous research on ant diversity documented variable responses to altitudinal gradients, which showed a certain dependency on macroclimate. For instance, studies reporting monotonic decreases in species richness, were frequent in temperate mountains [19–21], while monotonic increases were observed in arid environments [22, 23]. Longino and Colwell [24] and Nowrouzi *et al.* [25] indicated more complex patterns in tropical climates, which often combined a monotonic decrease at high elevations with a plateau at low elevations. Mid-elevation peaks were reported in tropical areas [26, 27], although also frequent at other latitudes [28, 29]. These patterns were partially attributed to geometric constraints, such as the mid-domain effect and area availability [30, 31] or the Rapoport rescue effect [32]. However the role of climatic factors, including temperature [19–21] and water availability [22, 29, 33, 34] as direct drivers of ant diversity is also unequivocal. Moreover, several studies demonstrated these climatic variables also indirectly modulated ant diversity, for example, by affecting primary productivity [33, 35–37].

Understanding how ant diversity responds to altitudinal gradients is not sufficient, however, to predict how environmental changes (e.g. rising temperatures) will affect species geographic ranges, because knowledge on species composition patterns is also required. Species assemblages and their distributions along ecological and geographical gradients result from species tolerances to climatic (abiotic) factors and biotic interactions, which interplay with niche evolution to determine community composition [38–40]. Kodric-Brown and Brown [41] defined a structured community as one where the organization is not due to randomness [41], and all situations in which communities are not random can be explained by spatial turnover, nestedness, or combinations of both [42]. Nestedness of species assemblages occurs when site biotas with smaller species numbers are subsets of the biotas at richer sites [43, 44]. If environmental and habitat filtering and not interspecific competition is responsible for nestedness, most species can coexist. Therefore, under increasing temperatures, species could migrate or expand their distribution elsewhere from the species former range, without necessarily implying other resident species will be displaced by the newcomers. On the contrary, spatial turnover implies the replacement of some species by others, exhibiting gains and losses of species from location to location globally [45]. This is a sign that some species cannot coexist, because of different climatic requirements, competitive exclusion, or both. In general, global warming will extend insect species ranges to higher altitudes [46], therefore with spatial turnover patterns, species will migrate up an altitudinal gradient and those at higher altitudes might suffer a subsequent reduction in geographic distribution range, and even disappear from the area. However, Maguire *et al.* [47] acknowledged both processes and community patterns might vary with scale, and exhibit spatio-temporal variation, which raises concerns on our capacity to generalize the results based on local studies. Indeed, transferability of results is increasingly considered in analyzing the effects of global change on species distributions, although such efforts are rarer in studies of community patterns [48–50].

In the present study, we explored patterns of ant species diversity and community composition in Mediterranean mountain grasslands, integrating spatio-temporal transferability. Specifically, we addressed the following questions: (1) what is the relationship between ant species richness and elevation; (2) how does elevation affect ant species composition; (3) what are the relative contributions of spatial turnover and nestedness in ant community composition; and (4) are our findings transferable spatially and temporally?

## Methods

### Study areas and sampling design

We studied grassland ant communities in two mountains in the larger Sistema Central range, Sierra de Guadarrama (reaching 2428 m a.s.l. at its highest peak) and Serrota (maximum 2294 m a.s.l.). The two mountains are ~ 100 km apart with similar climatic, geologic, and biotic characteristics. Mean annual temperatures range from ~ 14 °C at 600 m to ~ 4 °C at the summits (2300 - 2428 m); and mean annual rainfall ranges from 550 mm to 1500 mm, with severe summer drought [51]. Substrata are primarily composed of granites, and pasturelands are distributed along the complete elevational gradient, largely as the result of traditional livestock grazing.

Sample sites were interspersed along the elevational gradient, facing southeast to southwest with gentle slopes (< 5%), and not affected by anthropogenic disturbance (i.e., not close to buildings or main paths). We sampled 18 grasslands in 2014 (in Guadarrama range) and 12 in 2015 (six in Guadarrama range and six in Serrota range). On each grassland, we randomly placed a 4 × 3 grid of pitfall traps, 2.5 cm diameter × 5 cm deep, with 5 m spacing between traps, which were filled with a 3:1 ethanol/ monoethylene glycol mixture. Ant sampling was conducted in July, during stable anticyclonic meteorological periods (Guadarrama: 17 - 23 July 2014 and 9 - 16 July 2015; Serrota: 1 - 8 July 2015), taking advantage of the seasonal ant activity peak during summer in Central Spain [52]. Traps were collected after one week and ant specimens were sorted to species, excluding winged ones.

Pitfall trapping is considered more objective and unbiased than other methods for sampling ground ants [53, 11]. We chose pitfall traps previously tested in other studies in Central Spain, which demonstrated success at capturing the complete species pool and the entire ant size range variability present in the sampled area [52, 54, 55].

### Statistical analyses

We examined the relationship between species richness, taxonomic composition, and nestedness of grassland ant communities with elevation. Our analytical strategy was to build descriptive models for the data gathered during the most intensive sampling in Guadarrama 2014, and to validate the patterns detected with data gathered in a subset of the Guadarrama grasslands during 2015 (temporal validation) and with data gathered in the Serrota range during 2015 (spatial validation).

#### Species richness

First, we assessed the relationship between species richness and elevation. We estimated the species pool size at each locality with the incidence-based first order jackknife estimator. The relationship shape between estimates of richness and elevation was explored applying a generalized additive model (GAM) with Gaussian errors and a thin plate regression spline for elevation. Models were built with data from Guadarrama 2014 sampling and validated with data from the Guadarrama 2015 and Serrota 2015 samplings. Squared Pearson correlation between jackknife estimates of richness and jackknife estimates predicted by the model were used to assess species richness pattern constancy between years and sites.

#### Taxonomic composition of assemblages

We explored how species assemblages differed based on taxonomic composition by applying non-metric multidimensional scaling (NMDS) to the hemi-matrices of binary Bray-Curtis distances among grasslands derived from species occurrences in pitfall traps. In addition, a model-based analysis was conducted by building separate negative binomial regressions for each species abundance data (as the multivariate response variable) by elevation, year, and interaction (as the explanatory variables) [56]. Significance of the interaction (assessed by resampling) would suggest that patterns of species composition along an elevational gradient changed between years. Likewise, for Guadarrama 2014 and Serrota 2015, a multivariate model was built by fitting negative binomial regressions for individual species abundance, elevation, location, and interaction [56]. Significance of the interaction would suggest how assemblages changed with elevation were context-dependent.

#### Nestedness patterns

To assess assemblage nestedness among grasslands surveyed on each mountain and year we calculated the matrix temperature (T) [57] and the NODF (Nestedness metric based on Overlap and Decreasing Fill) [58]. While T has traditionally been the most commonly used metric for assessing overall nestedness [59], the more recent NODF index is claimed to exhibit more robust statistical properties, and separate contributions to nestedness of columns (due to species incidence) and rows (due to site composition) can be quantified [58, 60]. T decreases and NODF increases with nestedness.

The significance of nestedness indices was estimated by comparison with suitable binary null models, which were simulated by randomizing the original matrix transformed to presence/absence binary data. Two null models were built to encompass the range from maximally liberal (equitable) to maximally conservative (constrained) null models [61]. The first was an equiprobable model built by randomizing rows and columns and therefore maintaining only the number of species presences (‘r00’ algorithm). The second was a proportional resampling model constructed by constraining randomization to maintain row and column totals (site richness and species incidence, respectively), while using marginal column frequency to select species (‘quasiswap’ algorithm). Thus, the first null model was more liberal and only accounted for matrix fill (the incidence of the total species set), while the second null model was more stringent, because it imposed additional structure on the data, accounting for among-site differences (e.g. different carrying capacities) and among-species differences (e.g. different rarities). Subsequently, significant nestedness was attributed to variation in observed species richness or species incidence, if evaluated with the equiprobable model, or to variation beyond that observed, if evaluated with the proportional resampling model [62]. One thousand randomizations of the original matrix were applied.

The relative degree of nestedness, i.e. how much a grassland was nested within the set of sampled grasslands in the same geographic area and time period, was evaluated by a nestedness rank, according to T. Poorer grasslands, i.e. those having lower species diversity which were a subset of richer grasslands, were assigned higher ranks. These ranks were estimated as the ordinate in nestedness plots built with the T index, which were calculated as (k - 0.5)/n for k = 1…, n rows (e.g. the bottom row in the graph, where k = 18 was the more nested site, which for the 2014 sampling in Guadarrama had a rank of (18 - 0.5)/18 = 0.97).

The shape of the relationship between nestedness and elevation during the 2014 sampling in Guadarrama (n = 18) was explored using GAM with Gaussian errors and a thin plate regression spline for elevation. This model was validated using Guadarrama 2015 data (temporal validation). Detection of a high correlation between observed (in 2015) and predicted ranks would suggest the nestedness rank of plots did not vary between years. Similarly, the model built with Guadarrama 2014 sampling data was validated using the Serrota 2015 data (spatial validation). Detection of a high correlation between observed and predicted ranks would suggest the nestedness rank of plots did not vary between regions.

The contributions of spatial turnover and nestedness to the distribution pattern in Guadarrama 2014 data were calculated using three beta diversity indices: Sørensen-based multiple-site dissimilarity (β_SOR_), Simpson-based multiple-site dissimilarity (β_SIM_), and nestedness-resultant multiple-site dissimilarity (β_NES_). β_SIM_ accounts for spatial turnover and β_NES_ integrates dissimilarity due to nestedness, while β_SOR_ expresses the total dissimilarity between communities and equals the sum of β_SIM_ and β_NES_ [42]. *P*-values were estimated through the equiprobable (‘r00’) null model (the proportional resampling null model was not useful to assess this partitioning of beta diversity).

Statistical analyses were performed using R (v 3.3.2; R Core Team 2016) and specialized packages vegan (v. 2.4-5) [63], mvabund (v. 3.12-3) [56], and betapart (1.5.0) [64].

## Results

We detected 37 species, all recorded in Guadarrama (35 in 2014 and 26 in 2015, 24 were shared between years) and 20 species in Serrota (all species shared with Guadarrama). A total of 15 species were common among the three surveys (**S1 Table**).

Jackknife estimates of ant species richness in Guadarrama 2014 ranged from three (SE = 0) species in most of the highest grasslands to 18.5 (SE = 2.6) at intermediate elevations (Table 1). Estimates of richness related to elevation were positive below 1100 m a.s.l. and negative until a lower limit was reached at ca. 2000 m a.s.l. (Fig 1). Overall, richness exhibited a hump-shaped relationship with elevation, which we described using GAM with a spline for elevation, roughly equivalent to a third-order degree polynomial (equivalent df = 3.96, F = 12.94, *P* < 0.0001, D^2^ = 83.8%, Fig 1). This pattern of species richness was similar between years and between mountain ranges (correlations for estimates of richness and richness predicted by the model were r = 0.92, t = 4.68, df = 4, *P* = 0.0095, R^2^ = 85% for Guadarrama 2015 data; and r = 0.86, t = 3.42, df = 4, *P* = 0.0267, R^2^ = 75% for Serrota 2015 data).

**Table 1.** Observed (S) and first-order jackknife estimator (J ± SE) of ant species richness in dry grasslands in central Spain.

| ID | Sampling | S | $J \pm SE$ | N | X | Y | Elevation (m) |
| --- | --- | --- | --- | --- | --- | --- | --- |
| G1 | G2014 | 7 | $7 \pm 0$ | 13 | 437876 | 4494480 | 685 |
| G1 | G2015 | 7 | $7.9 \pm 0.9$ | 10 | 437876 | 4494480 | 685 |
| G2 | G2014 | 8 | 8±0 | 12 | 434821 | 4496549 | 768 |
| S1 | S2015 | 14 | 18.5±2.4 | 10 | 351711 | 4474792 | 808 |
| G3 | G2014 | 14 | 16.7±1.6 | 11 | 440394 | 4503515 | 865 |
| G4 | G2014 | 14 | 16.8±1.6 | 12 | 435541 | 4506094 | 1012 |
| G5 | G2014 | 14 | 15.8±1.3 | 12 | 431692 | 4513777 | 1044 |
| G5 | G2015 | 11 | 11.9±0.9 | 12 | 431692 | 4513777 | 1044 |
| S2 | S2015 | 11 | 14.7±2.3 | 12 | 357653 | 4481333 | 1086 |
| G6 | G2014 | 13 | 14.8±1.3 | 9 | 414145 | 4507982 | 1171 |
| G7 | G2014 | 13 | 18.5±2.6 | 12 | 438808 | 4523233 | 1285 |
| G8 | G2014 | 15 | 15.9±0.9 | 11 | 416330 | 4511347 | 1330 |
| G8 | G2015 | 14 | 17.5±2.2 | 8 | 416330 | 4511347 | 1330 |
| S3 | S2015 | 13 | 15.8±1.6 | 12 | 340207 | 4476196 | 1374 |
| G9 | G2014 | 9 | 9±0 | 12 | 436806 | 4524537 | 1487 |
| G10 | G2014 | 10 | 12.8±1.6 | 12 | 400821 | 4495703 | 1625 |
| S4 | S2015 | 4 | 4±0 | 11 | 326465 | 4482790 | 1646 |
| G11 | G2014 | 3 | 3±0 | 19 | 400543 | 4496999 | 1675 |
| G12 | G2014 | 5 | 5.9±0.9 | 12 | 419833 | 4519838 | 1786 |
| G12 | G2015 | 5 | 5.9±0.9 | 12 | 419833 | 4519838 | 1786 |
| G13 | G2014 | 5 | 5±0 | 12 | 419076 | 4519012 | 1830 |
| G14 | G2014 | 4 | 4.9±0.9 | 12 | 418737 | 4519177 | 1920 |
| S5 | S2015 | 2 | 2±0 | 12 | 323719 | 4482788 | 2000 |
| G15 | G2014 | 3 | 3±0 | 12 | 419392 | 4520829 | 2026 |
| G15 | G2015 | 3 | 3±0 | 8 | 419392 | 4520829 | 2026 |
| G16 | G2014 | 3 | 3.9±0.9 | 12 | 417883 | 4521249 | 2139 |
| G17 | G2014 | 4 | 5.8±1.9 | 12 | 418676 | 4521128 | 2266 |
| S6 | S2015 | 3 | 3.9±0.9 | 12 | 323944 | 4485467 | 2286 |
| G18 | G2014 | 3 | 3±0 | 12 | 419420 | 4522544 | 2390 |
| G18 | G2015 | 3 | 3.9±0.9 | 12 | 419420 | 4522544 | 2390 |

**Fig 1.**
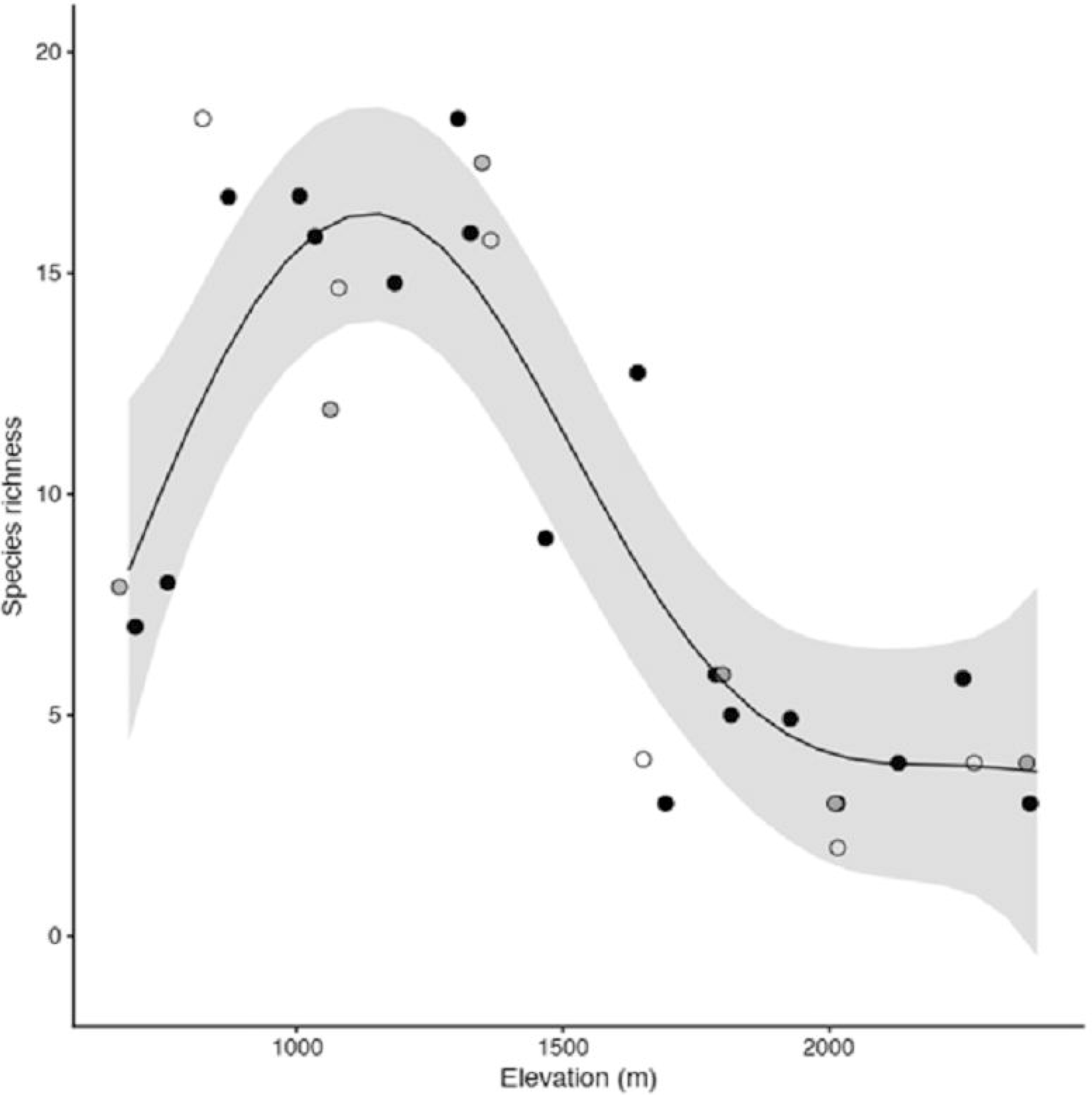
Relationship between first-order jackknife estimates of ant species richness (see Table 1) and elevation in dry grasslands of a Mediterranean range in central Spain.

Grassland plots were ordered following elevation (in m a.s.l.), identified with an ID, where G stands for Guadarrama, S stands for Serrota, and numbers represent grassland position in increasing order of elevation. Sampling: data from Guadarrama range in 2014 (G2014) and 2015 (G2015) and Serrota range in 2015 (S2015). N: samples sizes (number of pitfall traps recovered from each locality). Geographic coordinates (X and were provided as UTM European Datum 1950.

Curve and 95% IC from GAM with Gaussian errors fit to the Guadarrama 2014 survey (black circles). Richness estimates for the surveys in Guadarrama 2015 (grey circles) and Serrota 2015 (open circles) were superimposed. Elevation values were slightly jittered to avoid overlap among some points.

A NMDS solution was built that satisfactorily summarized the Bray-Curtis distances among grasslands in both years and study regions (stress = 0.10, linear R^2^ between Bray-Curtis and NMDS distances = 0.95). This scaling separated grasslands roughly according to elevation, but not regions or years and suggested an elevation threshold of 1700 m a.s.l. was a suitable elevation to distinguish the two groups (Fig 2). Multivariate GLMs for species abundances revealed significant elevation effects (*P* = 0.001) and a significant region effect (*P* = 0.047), but significant effects were not detected for year or any elevation interactions (Table 2). These results suggested taxonomic composition varied mainly with elevation, but differences between regions cannot be excluded.

**Table 2.** Multivariate GLMs (negative binomial regressions) fitted to species abundances. *P*-values estimated by bootstrap (n = 1000). Significant *P*-values (*P* < 0.05) are highlighted in bold. a) Guadarrama 2014 vs. Guadarrama 2015. b) Guadarrama 2014 vs. Serrota 2015.

|  | Deviance | P values |
| --- | --- | --- |
| a) |  |  |
| Elevation | 192.91 | **0.001** |
| Year | 50.27 | 0.649 |
| Elevation x Year | 5.99 | 0.997 |
| b) |  |  |
| Elevation | 178.14 | **0.001** |
| Region | 84.44 | **0.047** |
| Elevation x Region | 18.85 | 0.618 |

**Fig 2.**
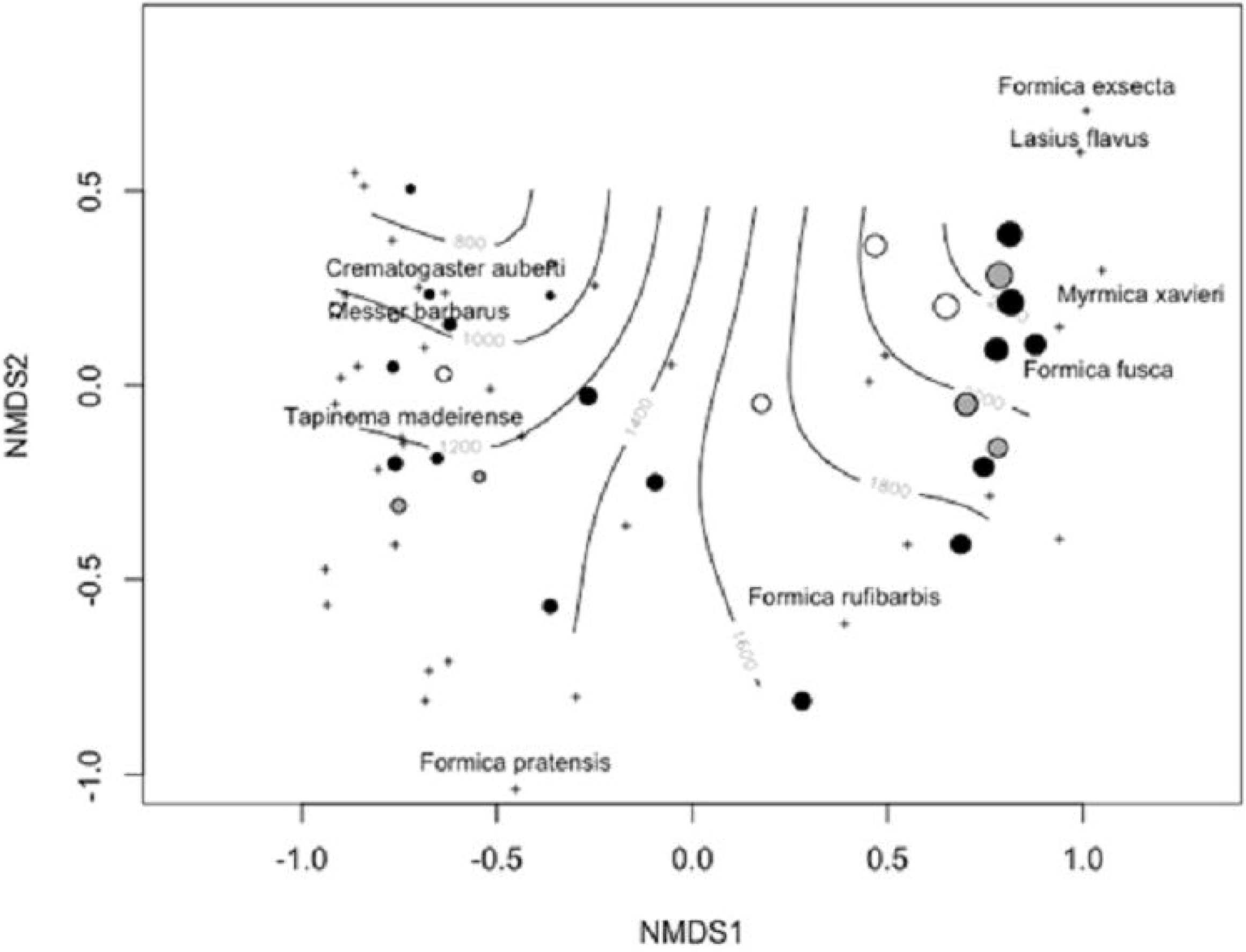
Non-metric multidimensional scaling (NMDS) of ant assemblages based on binary Bray-Curtis distances among grasslands calculated with species occurrences from pitfall traps in a Mediterranean range of central Spain.

Circles represent data from Guadarrama (black: year 2014, grey: year 2015) and Serrota (open circles) ranges. Symbol sizes are proportional to the elevation of sampling plots. As reference, circles are superimposed to a smooth elevation surface (estimated using a GAM of elevation on a bivariate spline of NMDS scores). The positions of some illustrative species are given as reference.

Ant species assemblages were nested, showing a pattern suggesting differences in species richness among localities were more important than differences in prevalence among species (nestedness among rows was higher than nestedness among columns, Table 3, **S2 Table**). Nestedness ranks for Guadarrama 2014 was described with GAM and curvilinear term (spline) for elevation, roughly similar to a third-degree cubic polynomial (df = 3.89, F = 15.38, *P* < 0.0001, D^2^ = 85.6%, Fig 3). The subset of these grasslands sampled in 2015 maintained their nestedness ranks and the model transferred to 2015 data (Pearson’s correlation between predicted and observed ranks, r = 0.96, t = 7.28, df = 4, *P* = 0.0018, R^2^ = 93%, **S3 Fig**). The model did not transfer between study regions (r = 0. 74, t = 2.18, df = 4, *P* = 0.10, R^2^ = 54%, **S4 Fig**). Largely, nestedness increased with elevation, with the exception of the lowest and highest elevation grasslands.

**Table 3.**
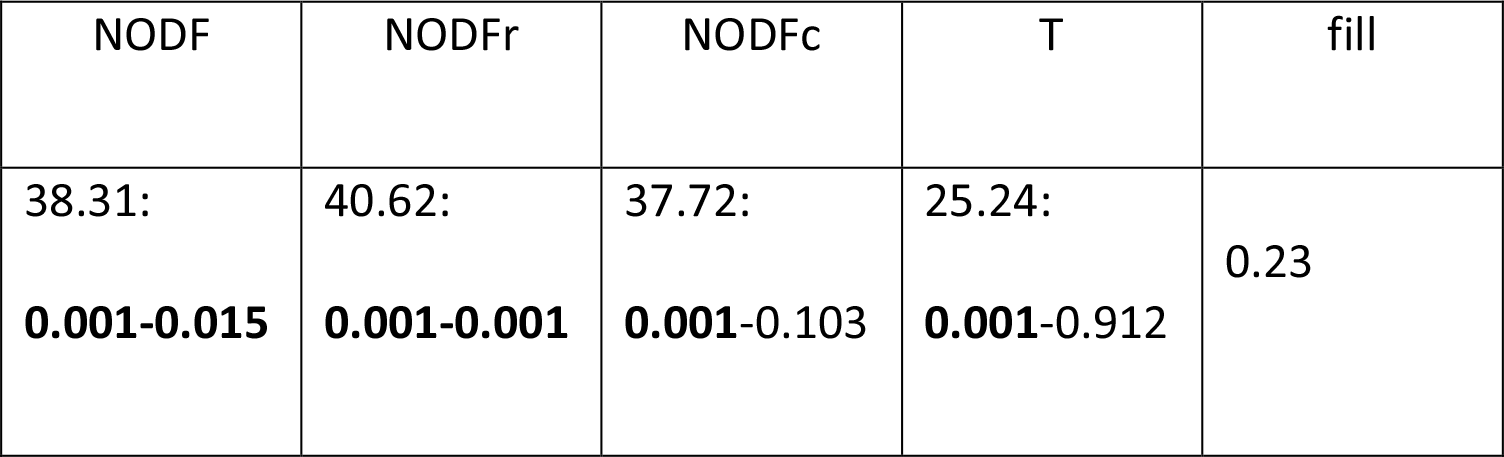
Nestedness indices for dry grassland ant assemblages in Guadarrama range 2014 surveys (18 grasslands, 35 species)

| NODF | NODFr | NODFc | T | fill |
| --- | --- | --- | --- | --- |
| 38.31:<br>**0.001-0.015** | 40.62:<br>**0.001-0.001** | 37.72:<br>**0.001-0.103** | 25.24:<br>**0.001-0.912** | 0.23 |

**Fig 3.**
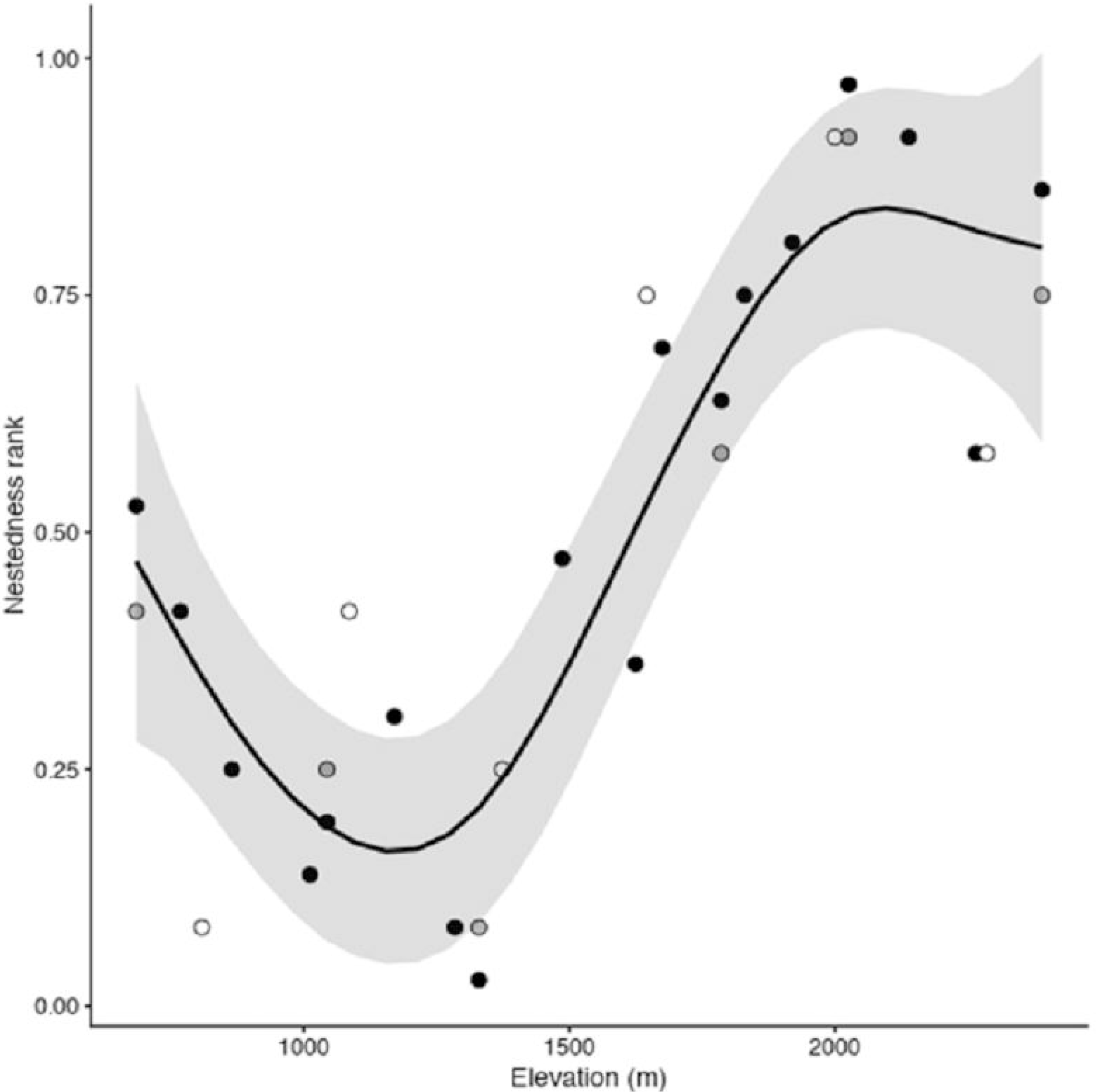
Generalized additive model (GAM) of nestedness ranks (based on nestedness temperature (T) index) for elevation in ant species assemblages in dry grasslands of a Mediterranean range in central Spain.

Indices were derived from the Nestedness metric based on Overlap and Decreasing Fill (NODF), NODFr for rows (i.e. localities) and NODFc for columns (i.e. species) and matrix temperature (T). Numbers under headings for indices provide the index values followed by *P*-values estimated by equiprobable (‘r00’) and proportional resampling (‘quasiswap’) binary null models. Significant *P*-values (*P* < 0.05) are in bold face. Fill: matrix fill (sum of 1s/sum of cells).

Curve and 95% IC from GAM with Gaussian errors fit to the Guadarrama 2014 survey (black circles). Nestedness ranks for the Guadarrama 2015 (grey circles) and Serrota 2015 (open circles) surveys are superimposed. Low ranks indicate poorer species community richness, where species are a subset of richer species communities. Elevation values were slightly jittered to avoid overlap among some points.

Species turnover and nestedness were significant beta diversity components, although species turnover among grasslands was more important than pure nestedness of assemblages (β_SOR_ = 0.88, β_SIM_ = 0.79, β_NES_ = 0.09, all *P*-values = 0.001). This relationship among beta diversity components did not vary among surveys (**S5 Table**).

## Discussion

Ant species richness and taxonomic composition changed predictably with elevation in the Mediterranean mountains we investigated, exhibiting largely consistent patterns between years and among regions. These changes were more notable along mid-elevations and associated more clearly with differences of species richness among sites than with prevalence or abundance among species. To our knowledge, this is the first study specifically designed to describe ant community species richness and taxonomic composition responses to elevational changes in the Mediterranean Basin. However, we restricted our study to grasslands, therefore the results might not be a suitable model system for other habitat types. Notwithstanding, grasslands are among the ecosystems exhibiting greater economic and ecological relevance in the Mediterranean Basin [65, 66].

Species richness showed a hump-shaped relationship with elevation, peaking at ca. 1100 m a.s.l. This model depiction fit to the primary dataset (Guadarrama 2014), transferred adequately to the 2015 Guadarrama dataset, and also the spatial validation dataset in the 2015 Serrota range. Despite geometric constraints as mid-domain effects were described to explain these curve types [30], more recent studies tended to downplay the importance of these mechanisms [24, 29, 67]. In fact, while mid elevation peaks in species diversity were reported as relatively common [28, 29], these observations are far from universal, suggesting regional factors play an important role in driving elevational patterns of species diversity. Interestingly, research showed monotonic increases in ant species richness were common in temperate climates [19–21], where elevation was correlated with temperature. In arid climates, monotonic decreases were observed [22, 23], where hydric stress was ameliorated with altitude. Indeed, hydric stress and temperature are the two main elevational correlates in Mediterranean climates, with water availability limiting communities at low elevations, while at high elevations cold temperature stress is more important [6, 7]. This double constraint might drive a hump-shaped spatial pattern of ant species richness in Mediterranean mountains.

Most studies on ant communities reported species richness positively responded to temperature [19–21, 24, 26, 27, 67, 68], but a number of findings also emphasized the role of water availability [22, 29, 33, 34, 69]. Cold temperatures can reduce local ant species richness by direct effects that limit ant activity, such as reduced foraging time [35, 70], and, at larger biogeographical scales, cold temperatures can also limit species pools by lowering speciation rates [67]. Azcárate *et al.* [70] reported low humidity, in turn, can limit colony activation by reducing foraging possibilities, and therefore constitutes a main environmental filter. Alternatively, temperature and water availability can facilitate ant species richness indirectly, these abiotic factors affecting primary productivity and therefore the range and abundance of resources ant colonies exploit [33, 35–37]. Diversity-productivity hypothesis states that as productivity increases, so does the availability of energy and resources, so density and size of colonies also increase. These relationships should increase local species richness by abundance, lowering local extinction risks or activating a sampling mechanism [71, 72]. In Mediterranean climates, primary productivity can be limited by cold temperatures and summer drought [73], so maximum primary productivity values likely occur at intermediate elevations, where summer drought is not too severe and temperatures are not too cold. Therefore, if species richness increases with primary productivity, a humped curve is also expected.

As anticipated, elevation also affected the taxonomic composition of ant communities. We observed a ca. 1700 m threshold separating high mountain ant species communities from those along the lower elevational gradient. This threshold was consistent among regions and between years. Furthermore, although nestedness was a significant component of beta diversity, most elevational shifts among communities were attributable to species turnover. This result suggested the communities with lowest ant species diversity (high mountain communities) were also the most unique in a regional context. The singularity of high elevation ant communities was previously observed in different biomes [22, 24, 34, 35], which reinforced the conservation value of the Mediterranean mountains reported for other taxa [74, 75].

The examination of community nestedness suggested current ant assemblages are context-dependent and likely highly vulnerable to global warming and other anthropogenic changes. This is because, first, species turnover exhibited a dominant role over pure nestedness, which indicated different and characteristic groups of ant species occurred along an elevational gradient, and at the very least, ant species community composition was distinct at high elevations (> 1700 m) (Fig 2). We want to emphasize that if species distributions followed gradually changing abiotic conditions (e.g. a decreasing temperature gradient with elevation), then we would expect a concomitant change in the taxonomic composition of ant assemblages, rather than the development of new, characteristic, species groups [42, 76]. It is also illuminating to consider that nestedness arises with contributions from differences among species and among sites [59]. Nestedness patterns elsewhere were explained as a simple sampling mechanism, where the smaller species group in less diverse, i.e. poorer communities were a subset of species in more diverse, i.e. richer communities, reportedly when the studied localities differed in resource availability (or productivity) and this factor determined species richness [72, 77]. However, this implied species attributes notably contributed to nestedness, because rarer species with small distributional ranges (often specialists or poor-dispersers) were not generally sampled by the least diverse, i.e. poorest communities, while the more common taxa exhibited a wide distributional range [61]. In our study, the least diverse communities were more nested than diverse ant communities, but nestedness among species was less important than among sites. We conclude the sampling mechanism did not fully explain the nested patterns in the ant communities we identified and habitat and abiotic filtering provided more plausible explanations for the patterns. However, we carefully controlled by design the habitat type (arid grasslands in gentle south facing slopes), which implied those habitat filters – if an integral component – should be fine grained. Nestedness patterns along the elevational gradient were temporally constant, but varied between mountain ranges, indicating ant community composition was spatially dependent. Finally, an historic explanation of current ant species distribution patterns in a landscape context is an important element to include. We studied a mountain range within the Mediterranean biome where major landscape changes have not occurred over the last thirty years. Extensive afforestation was conducted during 1950’s and a progressive abandonment of the region took place between the 1960’s and 1980’s. Therefore, we assume all sites are equally accessible from the regional species pool. However, we cannot disregard other time-related mechanisms, which might explain community composition, such as priority effects favoring one or more species, depending on their relative order of arrival to a site [78], or metapopulation dynamics involving local extinctions and recolonizations in periods beyond this two-year study [79].

Overall, species taxonomic composition varied non-linearly with elevation. The insufficiency of the sampling mechanism emphasizes that ant assemblages are highly vulnerable to global changes on climate and anthropogenic impacts, which might affect species at the habitat and/or community level. It is unlikely species’ range shifts that track environmental change will maintain current ant species community composition.

## Acknowledgements

This study could not have been carried out without the enthusiastic support of students at Department of Ecology UAM (Cristina Rota, Víctor Pascual, Jesús Miranda, Celia Santos, Rubén Altozano, Inés Cueva) and friends (Montse Reina, Esperanza Iranzo, Aimara Planillo, Carlos Pérez Carmona, Xavier Espadaler) who helped with field and laboratory work. The study received partial support by the Madrid’s Government research group network REMEDINAL3-CM (S-2013/MAE-2719) and is a contribution to project CGL2014-53789-R.

## Supporting information

**S1 Table. Species presence in each survey.**

**S2 Table. Nestedness indices for dry grassland ant assemblages in surveys from all study areas in central Spain.** Guadarrama range (-G2014- year 2014: 18 grasslands, 35 species; -G2015- year 2015: 6 grasslands, 26 species) and Serrota range (-S2015- year 2015: 6 grasslands, 20 species). Indices are Nestedness metric based on Overlap and Decreasing Fill (NODF), NODFr for rows (i.e., localities) and NODFc for columns (i.e., species) and matrix temperature (T). Numbers under headings for indices provide index values followed by *P*-values estimated by equiprobable (‘r00’) binary null models and proportional resampling (‘quasiswap’). Significant *P*-values (*P* < 0.05) are in bold face. Fill: matrix fill (sum of 1s/sum of cells).

**S3 Fig. Temporal validation of the relationship between nestedness and elevation.** Generalized additive model (GAM) of nestedness ranks (based on nestedness temperature index) on elevation in ant assemblages from Guadarrama range (central Spain) fit to Guadarrama 2014 data and validated with Guadarrama 2015 data.

**S4 Fig. Spatial validation of the relationship between nestedness and elevation.** Generalized additive model (GAM) of nestedness ranks (based on nestedness temperature index) on elevation in ant assemblages from Guadarrama range (central Spain) fit to Guadarrama 2014 data and validated with Serrota 2015 data.

**S5 Table. Multiple-site dissimilarities of beta diversity.** Multiple-site dissimilarities accounting for the spatial turnover (β_SIM_) and the nestedness components (β_NES_) of beta diversity, and sum of both values (β_SOR_) for dry grassland ant assemblages in all study areas in central Spain. G2014: Guadarrama range 2014; G2015: Guadarrama range 2015; S2015: Serrota range 2015. *P*-values estimated by equiprobable binary null models (‘r00’) are given between parentheses. Significant *P*-values (*P* < 0.05) are in bold face.

